# Host range projection of SARS-CoV-2: South Asia perspective

**DOI:** 10.1101/2020.09.30.320242

**Authors:** Rasel Ahmed, Rajnee Hasan, A.M.A.M Zonaed Siddiki, Md. Shahidul Islam

**Author notes:** Correspondence: **Md. Shahidul Islam**, Basic and Applied Research on Jute Project Bangladesh Jute Research Institute, Dhaka-1207, Bangladesh.

## Abstract

Severe Acute Respiratory Syndrome Coronavirus 2 (SARS-CoV-2), the causing agent of Coronavirus Disease-2019 (COVID-19), is likely to be originated from bat and transmitted through intermediate hosts. However, the immediate source species of SARS-CoV-2 has not yet been confirmed. Here, we used diversity analysis of the angiotensin I converting enzyme 2 (ACE2) that serves as cellular receptor for SARS-CoV-2 and transmembrane protease serine 2 (TMPRSS2), which has been proved to be utilized by SARS-CoV-2 for spike protein priming. We also simulated the structure of receptor binding domain of SARS-CoV-2 spike protein (SARS-CoV-2 S RBD) with the ACE2s to investigate their binding affinity to determine the potential intermediate animal hosts that could spread the SARS-CoV-2 virus to humans in South Asia. We identified cow, buffalo, goat and sheep, which are predominant species in the household farming system in South Asia that can potentially be infected by SARS-CoV-2. All the bird species studied along with rat and mouse were considered less potential to interact with SARS-CoV-2. The interaction interfaces of SARS-CoV-2 S RBD and ACE2 protein complex suggests pangolin as a potential intermediate host in SARS-CoV-2. Our results provide a valuable resource for the identification of potential hosts for SARS-CoV-2 in South Asia and henceforth reduce the opportunity for a future outbreak of COVID-19.

## INTRODUCTION

Coronavirus Disease 2019 (COVID□19) caused by Severe Acute Respiratory Syndrome Coronavirus 2 (SARS-CoV-2)^1,2^ has become a pandemic and spread ∼213 countries. The disease has already claimed more than 1 million lives and around 33 million infections worldwide as of 28 September, 2020 (https://www.worldometers.info/coronavirus). These figures are still growing in an alarming rate. The pandemic creates an overwhelming situation which frightens the public health and economy worldwide with an enormous effect.

The comparative analysis of SARS-CoV-2 sequence with other related viruses showed that bats are likely to be the originating species of this virus^3^. The zoonotic origin of SARS-CoV-2, predicted by some early cases which were linked to the Huanan seafood and wild animal market in Wuhan city of China, also raised the possibility of the transmission of SARS-CoV-2 through an intermediate host^4^. Severe Acute Respiratory Syndrome Coronavirus (SARS□CoV) and Middle East Respiratory Syndrome Coronavirus (MERS□CoV) were also originated from bat species and considered to be transmitted to human via intermediate host palm civets^5^ and dromedary camels^6^ respectively. However, the intermediate host or immediate source species of SARS-CoV-2 has not yet been confirmed. In the late 19^th^ century, there was a pandemic caused by a human coronavirus^7^ which might have originated from cattle or swine^8^. In addition, cross species transmission was reported in some studies where it was shown that coronaviruses can be transmitted from bats to human and other wildlife species^9^ and also from human to tiger^10^. Therefore, it is important to know the possible host range of SARS-CoV-2 for taking effective measures to control the pandemic and further cross species transmission. Detection of viruses from individuals of all the species present within a certain region in a pandemic situation is very difficult in terms of time and labor required for the lab work. Prediction of potential host species with in silico approaches based on the known factors of host-pathogen interaction could make the task feasible.

Host range determination and pathogenesis of coronaviruses largely depend on the receptor recognition by viral spike (S) protein^11^. Receptor□binding domain (RBD) of SARS-CoV-2 spike protein (SARS-CoV-2 S RBD) utilizes angiotensin□converting enzyme 2 (ACE2) as the functional receptor of the host to enter the cell^12^. Human ACE2 was identified as the cellular receptor for SARS-CoV also^13^. ACE2 is primarily associated to work in controlling blood pressure and to regulate the function of kidney and heart^14^. A recent study showed that ACE2 of civet, swine and Chinese horseshoe bat could be utilized by SARS-CoV-2 as entry receptor^3^. Binding of the surface unit, S1, of the S protein with cellular receptor facilitates viral attachment to the surface of target cells and the transmembrane unit, S2, is required to allow fusion of viral and cellular membranes^15^. However, coronaviruses require S protein priming, cleavage at the S1/S2 and the S2, to facilitate the binding with receptor for viral entry to host cell and so does the need of cellular protease^16,17^.

SARS□CoV was reported to use transmembrane protease serine 2 (TMPRSS2) for S protein priming^17,18,19^. A recent study showed that beside ACE2 as the receptor, SARS-CoV-2 also utilizes TMPRSS2 as cellular protease for viral entry into primary target cells and for viral spread in the infected host^20^. Therefore, comparative analysis of ACE2 and TMPRSS2 of target species could be served as the method for quick screening of potential host range.

In this study, we considered the subsequent analysis of ACE2 and TMPRSS2 to predict the potential as well as the intermediate host of SARS-CoV-2 among the selected species, which are mostly common in human habitation of South Asia (Bangladesh, Bhutan, India, Maldives, Nepal, Pakistan, and Sri Lanka). More than 1.8 billion people, about 24% of the world population, are living in South Asia^21^ and more than 0.1 million deaths (∼10% of global deaths) and around 7 million cases (∼ 20% of global cases) were reported in this part of the world to covid-19 pandemic as of 28 September, 2020 (https://www.worldometers.info/coronavirus). However, recent works have only considered ACE2 and did not highlight the species commonly present in South Asia to predict the host range of SARS-CoV-2^22,23,24,25^ in this region. Here, we made use of the ACE2 protein sequences of 34 species and TMPRSS2 of 30 species, subjected to be categorized in terms of their utilization by SARS-CoV-2 S protein through comparative analysis.

## METHODS

### Sequences used in the study

Protein sequences of ACE2 and TMPRSS2 were retrieved from either UniProt Knowledgebase or the NCBI GenBank Database. The amino acid (aa) sequences of ACE2 used in this study were from 34 species whereas 30 TMPRSS2 sequences were considered as the TMPRSS2 sequences of palm civet, intermediate horseshoe bat, Chinese rufous horseshoe bat and Pearson’s horseshoe bat were not available in the databases. The species names along with the sequence IDs are given in Table 1.

**Table 1.**
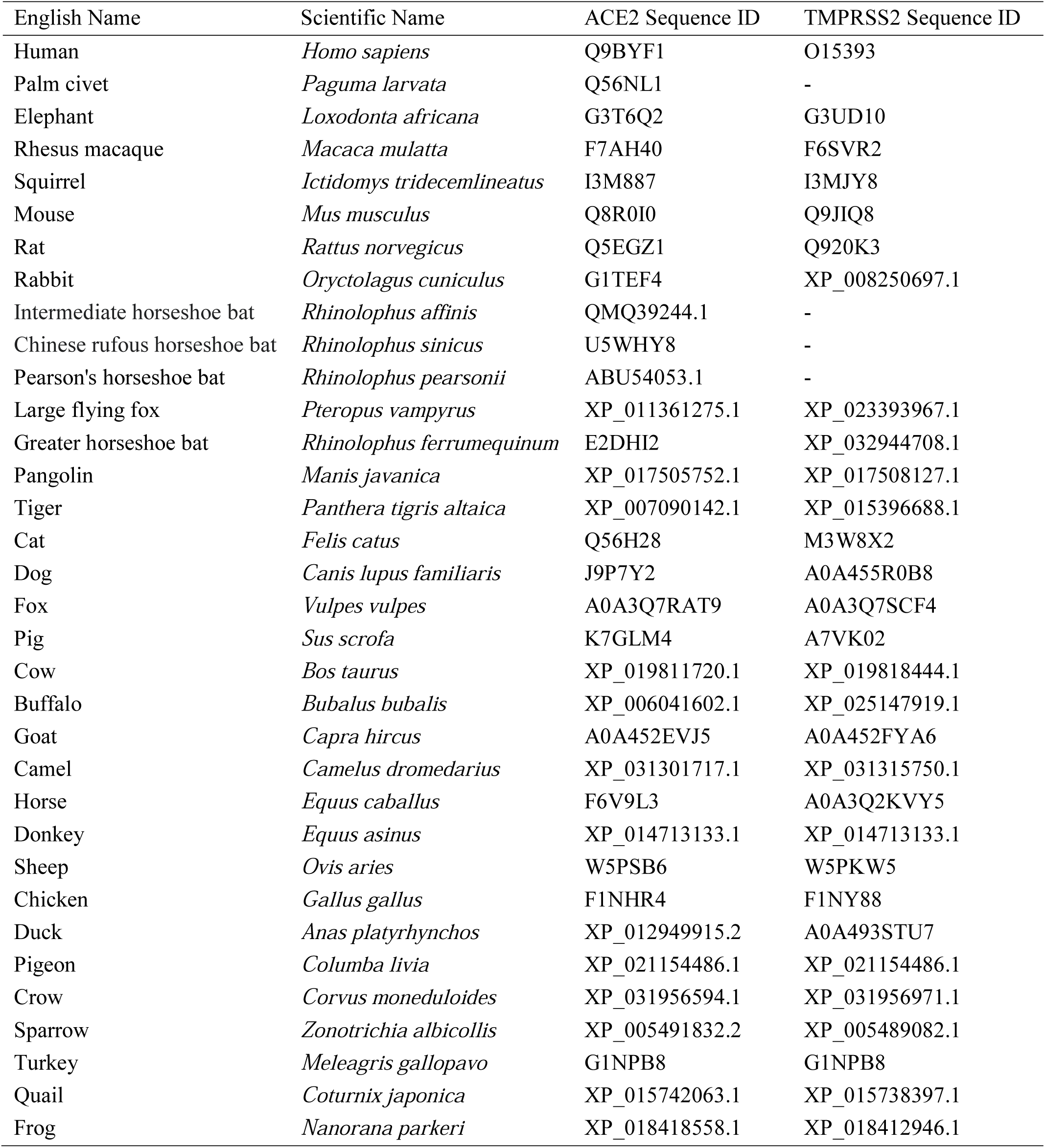
List of the species with sequence IDs

### Phylogenetic analysis

Evolutionary analysis was performed using molecular evolutionary genetics analysis version X (MEGA□X)^26^. Multiple comparisons were conducted using ClustalW multiple sequence alignment program incorporated in MEGA□X. Phylogenetic trees were generated with Jones□Taylor□Thornton evolutionary model using a maximum□likelihood method with 1,000 bootstrap replicates.

### SARS-CoV-2 S RBD and ACE2 binding capacity prediction

The critical aa sites for the utilization by SARS-CoV-2 were selected based on the known interactions of ACE2 and SARS-CoV-2 residues. A total of 20 aa sites were selected as important according to two recently solved crystal structures of the SARS-CoV-2 S RBD in complex with human ACE2^12,27^. We developed a scoring system for predicting the likelihood of the SARS-CoV-2 binding to ACE2s of 33 species. The initial score for every aa in the 20 selected residue sites which are identical with human ACE2 was set to 5. The substitution on each critical site produced a score of 0. The aa substitutions reported to abolish or strongly inhibit the receptor binding with SARS-CoV resulted in a reduction of its mark to −5. The final score was calculated by summation of the value of each of the 20 critical aa residues. The binding capacity of ACE2s of different species were ranked based on the final score. The ACE2s of the species scored below 50 were considered less potential to be utilized by SARS-CoV-2 S RBD.

### Homology-based structure simulation of ACE2□RBD complex

The recently solved crystal structure of SARS-CoV-2 S RBD^12^ bound to the cell receptor ACE2 was used as template (PDB: 6M0J) for homology modeling to simulate the interaction interfaces of SARS-COV-2 S RBD and ACE2 of human, pangolin, intermediate horseshoe bat, cow, goat, cat and dog using MODELLER^28^ incorporated in Chimera software version 1.14. The structures were visualized and analyzed with Chimera software version 1.14^29^.

## RESULTS

### Phylogenetic analysis of ACE2 and TMPRSS2

To investigate the molecular basis of evolution we studied overall sequence variation of ACE2 and TMPRSS2 among the selected species. For this purpose, phylogenetic trees were constructed based on the aa sequences of ACE2 and TMPRSS2 (Figure 1). Results showed that human ACE2 was clustered with rhesus macaque but separated from mouse and rat ACE2 in both of the trees. The members of Bovidae family such as cow, buffalo, goat and sheep formed a cluster and all the bird species were clustered with chicken in ACE2 and TMPRSS2 trees.

**FIGURE 1.**
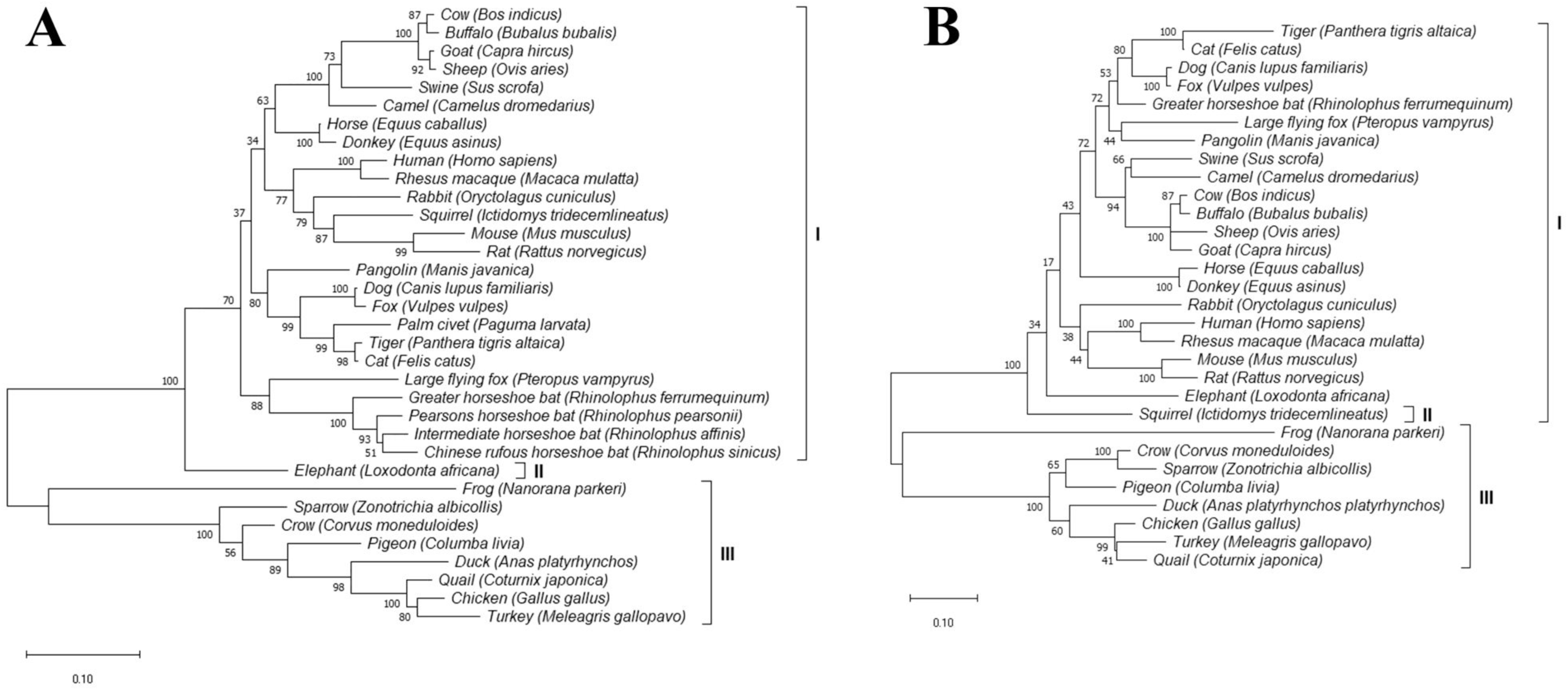
Phylogenetic analysis of protein sequences from the selected species. The trees were constructed on the whole aa sequences using maximum-likelihood method by MEGAX with 1,000 bootstrap replicates. A, phylogenetic tree of ACE2 protein sequences of different species. B, phylogenetic tree of TMPRSS2 protein sequences of different species.

### ACE2 utilizing capability by SARS-CoV-2 S RBD

We determined 20 potential aa residues of ACE2 important in interacting with SARS-CoV-2 based on the recently solved crystal structure of the SARS-CoV-2 S RBD in complex with human ACE2^12,27^. These aa residues were S19, Q24, T27, D30, K31, H34, E35, E37, D38, Y41, Q42, L45, L79, M82, Y83, E329, N330, K353, G354 and R393. The substitutions such as K31D, Y83F and K353H/A/D were reported to strongly inhibit or abolish the binding with SARS-CoV^30,31^. According to our scoring criteria where we set 5 marks for identical residue, 0 for substitution and −5 for substitution responsible for strong inhibition to bind with ACE2 compared to human ACE2 in corresponding site, we listed 34 species chronologically from highest to lowest score (Table 2). The utilization capacity of SARS-CoV-2 S RBD was predicted based on the cumulative final score achieved through summation of 20 individual scores. As shown in Table 2, only rhesus macaque achieved full score of 100 while squirrel got second highest score of 85. Species from the family Bovidae (cow, buffalo, goat and sheep) scored third highest of 80 along with cat, tiger and rabbit. Intermediate horseshoe bat scored 75 while other bats except greater horseshoe bat; pangolin, probable reservoir of SARS-CoV-2^32^; palm civet, considered as intermediate host of SARS-CoV ^5^; and camel, intermediate host of MERS-CoV^9^ scored between 60 to 70. According to our scoring system, the ACE2s of the species scored below 50 were considered less potential to be utilized by SARS-CoV-2 S RBD. All the bird species along with rat and mouse scored less than 50.

**Table 2.** Key aa comparison of ACE2 among different species interacting with SARS-CoV-2 S RBD

| Species | Position of aa |  |  |  |  |  |  |  |  |  |  |  |  |  |  |  |  |  |  |  | Score |
| --- | --- | --- | --- | --- | --- | --- | --- | --- | --- | --- | --- | --- | --- | --- | --- | --- | --- | --- | --- | --- | --- |
|  | 19 | 24 | 27 | 30 | 31 | 34 | 35 | 37 | 38 | 41 | 42 | 45 | 79 | 82 | 83 | 329 | 330 | 353 | 354 | 393 |  |
| Human | S | Q | T | D | K | H | E | E | D | Y | Q | L | L | M | Y | E | N | K | G | R | 100 |
| Rhesus macaque | S | Q | T | D | K | H | E | E | D | Y | Q | L | L | M | Y | E | N | K | G | R | 100 |
| Squirrel | S | L | T | D | K | Q | E | E | D | Y | Q | L | L | A | Y | E | N | K | G | R | 85 |
| Rabbit | S | L | T | E | K | Q | E | E | D | Y | Q | L | L | T | Y | E | N | K | G | R | 80 |
| Tiger | S | L | T | E | K | H | E | E | E | Y | Q | L | L | T | Y | E | N | K | G | R | 80 |
| Cat | S | L | T | E | K | H | E | E | E | Y | Q | L | L | T | Y | E | N | K | G | R | 80 |
| Cow | S | Q | T | E | K | H | E | E | D | Y | Q | L | M | T | Y | D | N | K | G | R | 80 |
| Buffalo | S | Q | T | E | K | H | E | E | D | Y | Q | L | M | T | Y | D | N | K | G | R | 80 |
| Goat | S | Q | T | E | K | H | E | E | D | Y | Q | L | M | T | Y | N | N | K | G | R | 80 |
| Sheep | S | Q | T | E | K | H | E | E | D | Y | Q | L | M | T | Y | D | N | K | G | R | 80 |
| Dog | S | L | T | E | K | Y | E | E | E | Y | Q | L | L | T | Y | E | N | K | G | R | 75 |
| Fox | S | L | T | E | K | Y | E | E | E | Y | Q | L | L | T | Y | E | N | K | G | R | 75 |
| Intermediate horseshoe bat | S | R | I | D | N | H | E | E | D | Y | Q | L | L | N | Y | N | N | K | G | R | 75 |
| Pig | S | L | T | E | K | L | E | E | D | Y | Q | L | I | T | Y | N | N | K | G | R | 70 |
| Camel | S | L | T | E | E | H | E | E | D | Y | Q | L | T | T | Y | D | N | K | G | R | 70 |
| Large flying fox | S | L | T | E | K | T | E | E | D | Y | Q | L | L | A | Y | E | K | K | G | K | 70 |
| Elephant | S | L | T | D | T | Q | E | E | D | Y | Q | L | L | D | F | E | N | K | G | R | 70 |
| Horse | S | L | T | E | K | S | E | E | E | H | Q | L | L | T | Y | E | N | K | G | R | 70 |
| Donkey | S | L | T | E | K | S | E | E | E | H | Q | L | L | T | Y | E | N | K | G | R | 70 |
| Pearson's horseshoe bat | S | R | T | D | K | H | E | E | D | H | E | L | L | D | Y | N | N | K | D | R | 70 |
| Pangolin | S | E | T | E | K | S | E | E | E | Y | Q | L | I | N | Y | E | N | K | H | R | 65 |
| Chinese rufous horseshoe bat | S | E | M | D | K | T | K | E | D | H | Q | L | L | N | Y | N | N | K | G | R | 65 |
| Palm civet | S | L | T | E | T | Y | E | Q | E | Y | Q | V | L | T | Y | E | N | K | G | R | 60 |
| Mouse | S | N | T | N | N | Q | E | E | D | Y | Q | L | T | S | F | A | N | H | G | R | 45 |
| Rat | S | K | S | N | K | Q | E | E | D | Y | Q | L | I | N | F | T | N | H | G | R | 45 |
| Greater horseshoe bat | S | L | K | D | D | S | E | E | N | H | Q | L | L | N | F | N | N | K | G | R | 45 |
| Chicken | D | - | T | A | E | V | R | E | D | Y | E | L | N | R | F | T | N | K | N | R | 35 |
| Pigeon | T | - | M | E | E | K | R | E | D | Y | E | L | - | - | - | N | N | K | N | R | 35 |
| Turkey | D | - | T | A | E | V | R | E | D | Y | E | L | N | R | F | T | N | K | N | R | 35 |
| Duck | D | - | M | A | E | V | R | E | D | Y | E | L | N | N | F | K | N | K | N | R | 30 |
| Crow | D | - | M | E | E | R | R | E | N | Y | E | L | N | S | F | E | N | K | N | R | 30 |
| Quail | D | - | K | A | E | V | R | E | D | Y | E | L | N | R | F | T | N | K | N | R | 30 |
| Sparrow | D | - | M | V | E | R | R | E | D | Y | E | L | N | K | F | N | N | K | N | R | 30 |
| Frog | D | D | V | Q | E | N | V | E | Q | H | Q | L | F | R | Y | N | K | L | D | R | 25 |

**Note:** Substituted residues compared to human ACE2 are colored with Green and substituted residues that were reported to strongly inhibit or abolish the binding with SARS-CoV-2 are colored with Red.

### Structure simulation by homology modeling of ACE2□RBD complex

The effect of residue substitutions at the atomic level is crucial for explaining the various receptor activities. To investigate such effects, we simulated the potential structures of SARS-COV-2 S RBD and ACE2 protein complex of human, pangolin, intermediate horseshoe bat, cow, goat, cat and dog using homology-based modeling based on the crystal structure of SARS-CoV-2 S RBD bound to the cell receptor ACE2 (PDB: 6M0J). Atoms of the substituted residues of ACE2s compared to corresponding residue of human ACE2 and atoms of the respective interacting residues of SARS-COV-2 S RBD in the binding interface were shown (Figure 2,3). We noticed some reduced or increased distances between the atoms of the interacting residues due to the substitutions. For instance, the atomic distances between N487 of SARS-COV-2 S RBD and corresponding substituted residues of ACE2 were 1.182Å, 2.626Å, 2.729Å and 3.218Å for pangolin (Figure 3A), intermediate horseshoe bat (Figure 3B), cat (Figure 3E) and dog (Figure 3F) respectively which was 2.013Å in case of human (Figure 2). Human ACE2 showed M as the aa in 82 position, while N82, N82, T82, T82, T82 and T81 was the corresponding aa in ACE2 of pangolin, intermediate horseshoe bat, cow, goat, cat and dog, respectively (Table 2). The atomic distance between the respective aa in 82 position of ACE2 and F486 of SARS-COV-2 S RBD was 3.052Å in human (Figure 2), 1.814Å in pangolin (Figure 3A), 2.102Å in intermediate horseshoe bat (Figure 3B), 2.679Å in cow (Figure 3C), 2.680Å in goat (Figure 3D), 2.856Å in cat (Figure 3E) and 2.575Å in dog (Figure 3F). In addition, the corresponding residue of G354 in pangolin was changed to H and this substitution produced more close contact with SARS-COV-2 (Figure 3A). Despite the aa changes, the interaction interfaces of these species ACE2s with SARS-COV-2 S RBD confirmed their potential interaction with this virus.

**FIGURE 2.**
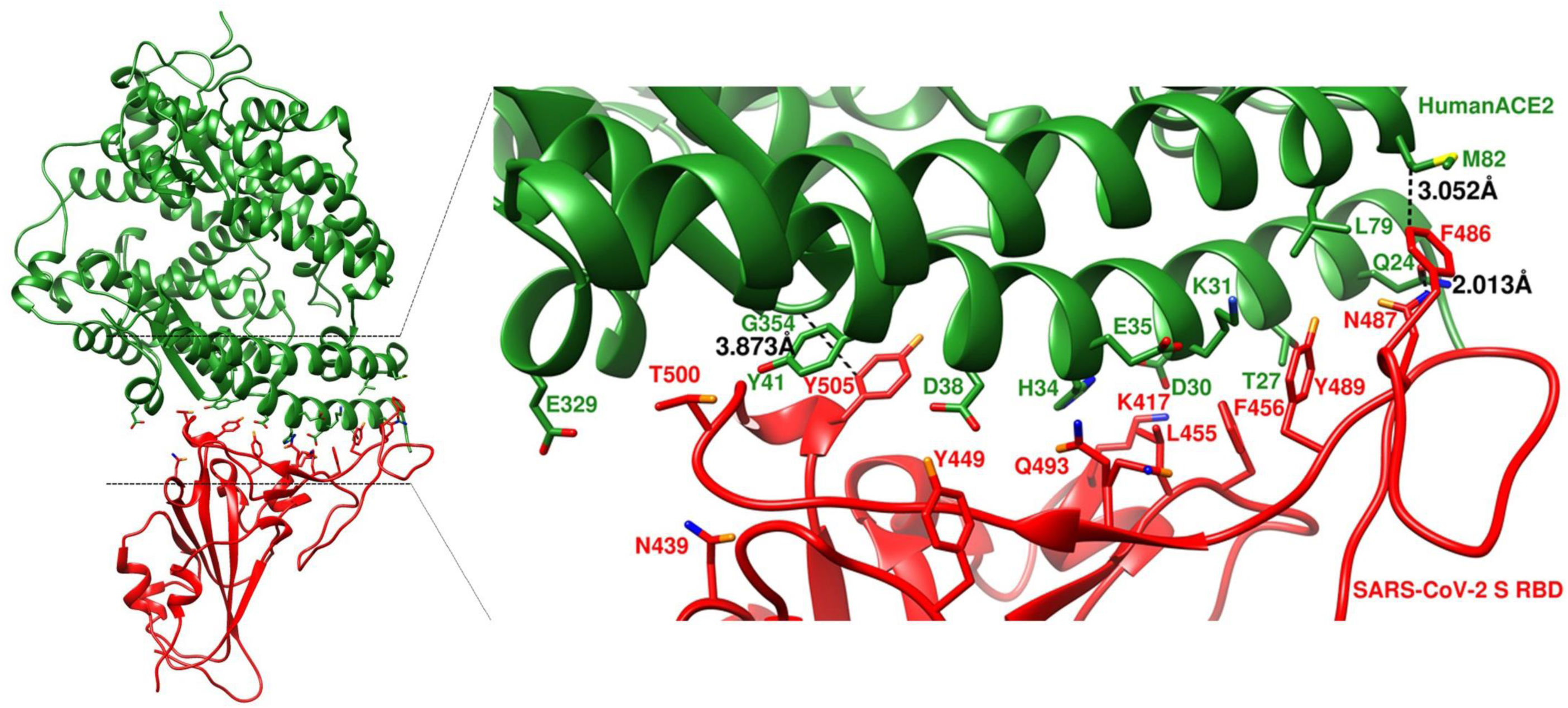
Overall structure simulation and structural details at the binding interface of SARS-COV-2 S RBD and human ACE2. HumanACE2 and SARS-COV-2 S RBD are in forest green and red, respectively.

**FIGURE 3.**
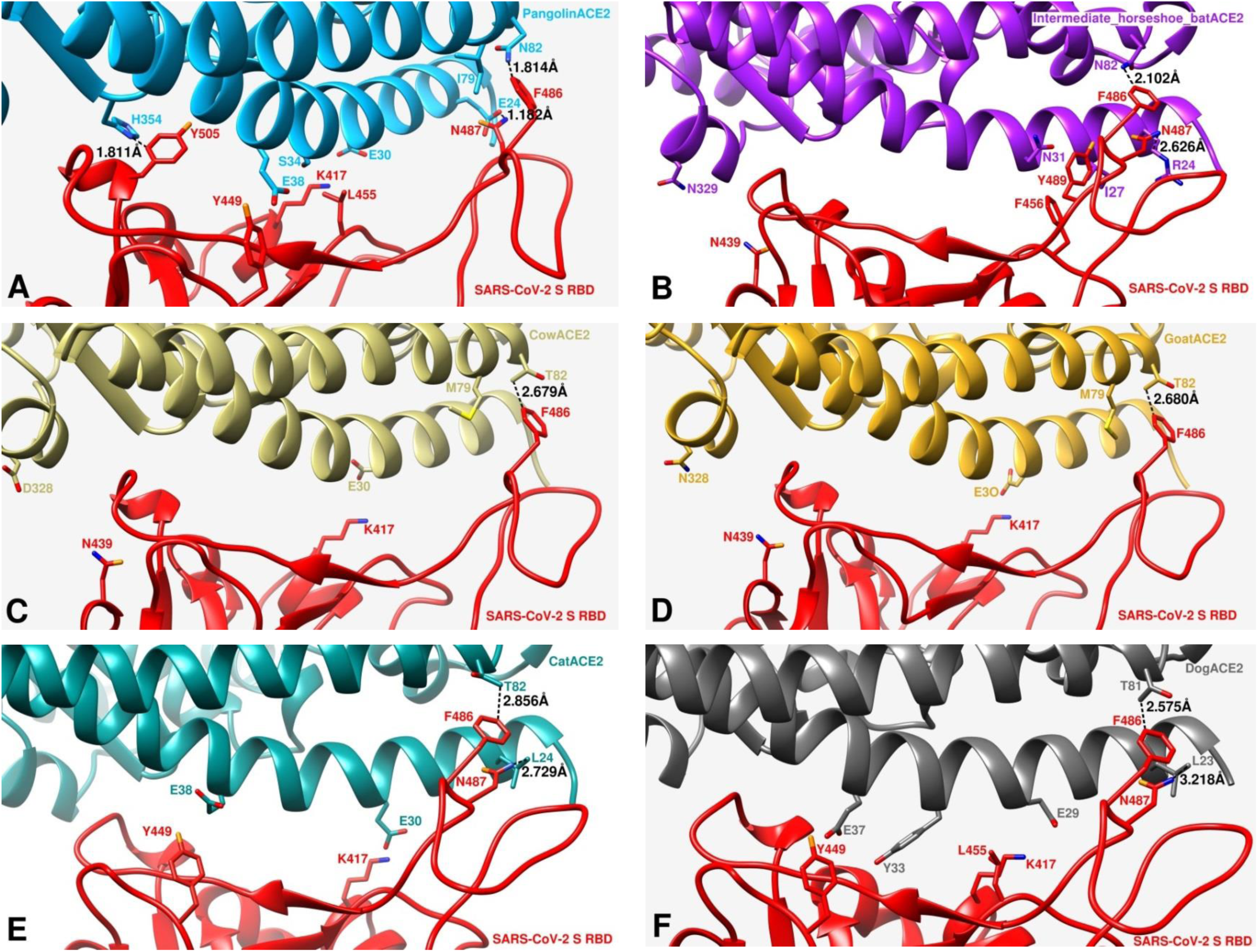
Comparisons of interactions at the SARS-CoV-2 S RBD and ACE2 interfaces. A, simulation of structural details at the binding interface of SARS-COV-2 S RBD and PangolinACE2. PangolinACE2 and SARS-CoV-2 S RBD are in deep skyblue and red, respectively. B, simulation of structural details at the binding interface of SARS-CoV-2 S RBD and Intermediate horseshoe batACE2. Intermediate horseshoe batACE2 and SARS-CoV-2 S RBD are in purple and red, respectively. C, simulation of structural details at the binding interface of SARS-CoV-2 S RBD and CowACE2. CowACE2 and SARS-CoV-2 S RBD are in dark khaki and red, respectively. D, simulation of structural details at the binding interface of SARS-CoV-2 S RBD and GoatACE2. GoatACE2 and SARS-CoV-2 S RBD are in goldenrod and red, respectively. E, simulation of structural details at the binding interface of SARS-CoV-2 S RBD and CatACE2. CatACE2 and SARS-CoV-2 S RBD are in dark cyan and red, respectively. F, simulation of structural details at the binding interface of SARS-CoV-2 S RBD and DogACE2. DogACE2 and SARS-CoV-2 S RBD are in dim gray and red, respectively.

## DISCUSSION

This is very important to determine the species range of a zoonotic disease like covid-19 to identify the potential intermediate host and natural reservoir of the causing agent for combating the disease in effective ways. Although the bat species, detected previously as the source of SARS-CoV^5^ and MERS-CoV^6^, was also identified as the probable origin of novel coronavirus SARS-CoV-2, the intermediate host of this virus is still not clear. We sought to identify the species, which have the probability to be infected with SARS-CoV-2 within the surroundings of human habitation in South Asia. For this purpose, we performed the diversity analysis of the selected species in terms on the aa composition of two proteins; ACE2, the host cell receptor of SARS-CoV-2^13^ and TMPRSS2, which has been proved to be utilized by SARS-CoV-2 for S protein priming^17,18,19^.

We generated phylogenetic trees to cluster the ACE2 and TMPRSS2 sequences of different species based on the evolutionary distance (Figure 1). Then, we looked for the residue level interaction of SARS-CoV-2 with these two proteins by analyzing protein complex structure. Unfortunately, the structure of TMPRSS2 or its interacting complex with SARS-CoV-2 is not reported yet. So, we took only ACE2 in consideration for further analysis. We intended to predict the capability of SARS-CoV-2 S RBD to utilize host cell receptor by investigating the substitution of critical aa in ACE2 of different species compared to human ACE2. We determined 20 aa critical in the binding interface of SARS-COV-2 S RBD and ACE2 protein complex according to two solved crystal structures^2,3^. We listed the species in descending order of respective score regarding the binding capacity of ACE2 with SARS-CoV-2 S RBD based on our scoring system (Table 2). Next, we tried to interpret the result of these two strategies to predict the likeliness of those species to interact with SARS-CoV-2.

Only the rhesus macaque which was reported to be as susceptible as human to SARS-CoV-2^33^, showed 100% similarity with human ACE2 (Table 2). Among the species which are very common in household farming system in South Asia-cow, buffalo, goat and sheep were predicted to be highly susceptible to SARS-CoV-2 by achieving the high score of 80. Special care should be taken to handle the management of these animals. Beside the bearing status as pet animal, a major population of cats and dogs are ownerless and seen frequently in the habitat of this part of Asia. According to the Table 2, cat gathered 5 points more than dog regarding the utilizing capability of ACE2 by SARS-CoV-2. This result is consistent with the study where SARS-CoV-2 was replicated efficiently in cat but poorly in dog^34^. Tiger ACE2 was clustered with cat (Figure 1A) and produced same score (Table 2). This finding is in agreement with the incident of a captive Malayan tiger which was reported to be positive with SARS-CoV-2 infection^10^. According to our findings, ACE2 of squirrel is an efficient (Table 2) receptor with second highest score of 85 suggesting its more susceptibility to SARS-CoV-2 infection than cat. Interestingly the intermediate horseshoe bat, whose affiliated coronavirus RaTG13 was reported to be most closely related with SARS-CoV-2 in terms of evolution, scored 75 which is the highest among the studied bat species. A range of the moderate score 60-70 was gained by other bat species and probable reservoir or intermediate host of SARS-CoV-2 (pangolin)^32^, SARS-CoV (palm civet)^5^ and MERS-CoV (camel)^6^. Our results differ with the findings where the animals such as civet^31^, Chinese rufous horseshoe bat^31^, pangolin^23^ and horse^23^ were predicted as the potential hosts with high risk for SARS-CoV-2 infection. Only the greater horseshoe bat within the bat species was found less potential. In addition, palm civet showing unique substitutions 37EQ and L45V and the large flying fox showing rare substitutions N330K and R393K (Table 2), could be the subject of further study to find the consequences of these alterations.

The correlation between genetic distances depicted by Figure 1 and scoring based ranking for interaction with SARS-CoV-2 (Table 2) was found interesting. The bird species along with frog, with scores less than 50% compared to human ACE2 (Table 2), could be ruled out from probable host range of SARS-CoV-2 and their clustering under group III in ACE2 tree (Figure 1A) further supported the idea. Despite achieving low score of 45, mouse, rat and greater horseshoe bat were clustered in group I where all the other studied species which produced substantial scores were clustered. Looking at the Table 2 we can see two changes; K31D in greater horseshoe bat and K353H in rat and mouse ACE2s. In addition, Y83F substitution was observed in these three species. All these alterations inhibit or abolish the binding of ACE2 with SARS-CoV^30,31^ and might be the reason of their inconsistent positioning in phylogenetic tree and scoring based ranking table. Our finding is also in agreement with the infectivity studies of rat and mouse^3,20^. Elephant was the only member of group II. It should be noted that, among the species with substantial scores only elephant bears inhibiting mutations in 83 AA position (Table 2). The overall clustering pattern of the species in TMPRSS2 tree (Figure 1B) is found almost similar to that of ACE tree (Figure 1A). Only exception is the position of squirrel and elephant. Compared to ACE tree, elephant was included to group I and squirrel was placed in group II as the only member of this group in TMPRSS2 tree. Although we could not predict the residue level utilizing capacity of TMPRSS2 by SARS-CoV-2 due to the unavailability of TMPRSS2 structure, from the above results we could say that the interaction pattern of TMPRSS2 with SARS-CoV-2 for the studied species would follow same prediction we inferred for ACE2 by analyzing Table 2 and Figure 1A. Beside in-depth analysis of ACE2, TMPRSS2 tree thus brought us extra confidence to our effort to predict potential hosts.

We further tried to investigate the binding affinity of ACE2 of six species (pangolin, intermediate horseshoe bat, cow, goat, cat and dog) along with human ACE2 with SARS-CoV-2 by simulating the potential structure of ACE2 and SARS-CoV-2 S RBD protein complex (Figure 2 and Figure 3) based on the recently solved crystal structure (PDB: 6M0J)^12^. We noticed that some residue changes in different ACE2 compared to human ACE2 may affect their interaction with SARS-CoV-2 S RBD in terms of the atomic distances within the interacting residues in the binding interface. Only the substitution in the 82 aa position is commonly seen to all the six species-substituted by N in pangolin and bat and by T in other four species (Table 2). Although all these substitutions showed reduction in the atomic distances in contacting F486 of SARS-CoV-2 S RBD (Figure 3), pangolin and bat produced closer contact than other four species. Our result agrees with the study which also claimed that N82 in ACE2 is closer to SARS-CoV-2 S RBD than M82^24^. In case of pangolin Q24E showed less distant contact with N487 of SARS-CoV-2 S RBD (Figure 3A), while substitution of aa in 24 position by other than E resulted more distant contact than human ACE2 (Figure 2 and Figure 3). In addition, G354H which is unique to pangolin produced more than 50% closer contact with Y505 of SARS-CoV-2 S RBD. The hypothesis that pangolin was involved in the evolution of SARS-CoV-2 is further supported by these results.

In a nutshell, we sought to combine phylogenetic analysis, identification and marking of key amino acids of host proteins utilized by SARS-CoV-2 to predict the potential host range among the species which are closely associated with daily activities of the people of South Asia region. Although this is plausible to predict the susceptibility to SARS-CoV-2 infection based on overall similarity of receptor residues, a single aa polymorphism or the glycosylated residue can disrupt the binding of the receptor to the virus^35^. Moreover, this might not be straight forward to identify individual reservoir within the same species because of intraspecies variation. Therefore, direct laboratory data and epidemiological investigation are required to make confirmed call. Nonetheless, our predictions provide quick assessment ahead of time-consuming individual laboratory experiments to detect risk associated species and so to develop proactive measure accordingly in controlling further cross species transmission. Our findings from structure simulation suggested pangolin as a potential intermediate host in SARS-CoV-2 evolution. Results of our analysis may also help to design optimized ACE2 for host mediated research to study SARS-CoV-2 infection.

## AUTHOR CONTRIBUTIONS

RA conceived the work, analyzed the data and wrote the manuscript. RA and RH contributed to graphics processing. RH, AMAMZS and MSI contributed to critical reviewing and editing the manuscript.

